# Temporal dynamics of gene expression in heat-stressed *Caenorhabditis elegans*

**DOI:** 10.1101/135988

**Authors:** Katharina Jovic, Mark G. Sterken, Jacopo Grilli, Roel P. J. Bevers, Miriam Rodriguez, Joost A. G. Riksen, Stefano Allesina, Jan E. Kammenga, L. Basten Snoek

## Abstract

There is considerable insight into pathways and genes associated with heat-stress conditions. Most genes involved in stress response have been identified using mutant screens or gene knockdowns. Yet, there is limited understanding of the temporal dynamics of global gene expression in stressful environments. Here, we studied global gene expression profiles during 12 hours of heat stress in the nematode *C. elegans*. Using a high-resolution time series of increasing stress exposures, we found a distinct shift in gene expression patterns between 3-4 hours into the stress response, separating an initially highly dynamic phase from a later relatively stagnant phase. This turning point in expression dynamics coincided with a phenotypic turning point, as shown by a strong decrease in movement, survival and, progeny count in the days following the stress. Both detectable at transcriptional and phenotypic level, this study pin-points a relatively small time frame during heat stress at which enough damage is accumulated, making it impossible to recover the next few days.

## Introduction

Heat-stress results from an exposure to potentially harmful temperature conditions beyond the optimum range of an organism. An increase in the intracellular temperature can interfere with protein homeostasis, leading to an accumulation of misfolded proteins and protein aggregates [1]. Over the past few decades, detailed insights have been obtained about the molecular and genetic control of the cellular response to heat-stress. To avoid the detrimental effects of cytotoxic misfolded protein species and protein aggregates, multiple stress response systems have evolved as a first line of defence to maintain proteostasis, of which the highly-conserved heat-shock response (HSR) pathway is prominent [1,2].

The accumulation of misfolded proteins is also a hallmark of aging and age-related diseases, such as Alzheimer’s and Parkinson’s disease [3–5]. The connection between the processes involved in stress and aging is further substantiated by the fact that several components of the stress response pathways were found to function as regulators of lifespan [6–8]. For example, the evolutionary highly conserved transcription factor HSF-1 is a key component in the initiation of the HSR, as well as a regulator of lifespan [9]. Therefore, understanding how an organism perceives and handles heat-stress is fundamental for understanding the molecular mechanisms that underlie aging [4].

The nematode *Caenorhabditis elegans* is an established metazoan model for studying the effect of - and response to – heat stress *in vivo* [4,6,7,9]. One of the most widely studied responses in *C. elegans* is to acute heat stress, which can be easily applied by exposing the animal to temperatures between 33-37°C [9–11]. It was shown that *C. elegans* detects and responds to heat stress via transient receptor potential channels and a neuropeptide signaling pathway [12], and is capable of swiftly up-regulating a suite of protective proteins (mainly chaperones) to prevent protein denaturation and misfolding, a process which affects every aspect of the animals’ biology [4]. Short or mild stresses can be tolerated and can even protect individuals against future stresses [11]. However, *C. elegans* is limited in the number of chaperones that can be produced, moreover, chaperones, being proteins themselves, are also likely to denature after sustained heat stress.

The effects of heat stress on *C. elegans* are often quantified on a phenotypic level by recording complex traits such as survival rate, mobility, and reproduction [11,13,14]. Generally, the inflicted damage accumulates with increasing temperature and exposure time. For example, brood size decreases with moderate increases in temperature beyond the optimum [14,15], whereas a strong decrease in survival rates is only observed after prolonged exposures to heat stress [11,16]. At the level of the transcriptome, a heat shock induces a strong response. Genome wide gene expression analysis in *C. elegans* shows that a two hour exposure to 35°C affects genes associated with development, reproduction and metabolism [17]. Furthermore, an exposure of 30 minutes to 33°C already induced a massive global gene expression shift highly dependent on HSF-1, affecting genes associated with a wide range of functions such as cuticle structure, development, stress response, and metabolism [18].

Yet, there is limited understanding of the temporal dynamics of global gene expression patterns during heat stress. Given the range of phenotypic effects, it is to be expected that the transcriptional response during heat stress is highly dynamic. For example, the initial transcriptional response to heat shock probably does not resemble the transcriptome after a lethal exposure to heat stress. To gain more insight into the underlying dynamics of the stress response, we have generated a high-resolution time-series of transcriptomic and phenotypic data of *C. elegans* exposed to heat stress conditions at 35°C for 0-12h. Transcriptomic analysis revealed a global shift in expression dynamics occurring between 3 and 4 hours into the heat exposure. The shift marks the end of an initially highly dynamic transcriptional response to heat stress that plateaus at longer exposures. On a phenotypic level, longer exposures (> 4h) were associated with much lower chances of recovery in the four days following the stress. Therefore, this phenotypic turning point follows shortly after the transcriptional turning point, and is marked by a strong decrease in movement, survival, and progeny count.

## Results

### Transcriptional variation during prolonged heat stress

We first assessed the impact of heat stress durations on genome-wide expression levels. Wild type Bristol N2 populations were exposed to heat stress conditions at 35°C for increased exposure durations between 0.5-12 hours (Fig 1A). To find the main sources of variation during the transcriptional response to heat-shock, we used principal component analysis (PCA). The first two principal components (PCs) captured 77% (1^st^ 57%, 2^nd^ 20%) of the total variation (Fig 1B). The first PC sorted the time points in chronological order, showing that variation in gene expression between samples was largely due to the increasing length of heat exposure. Together, the 1^st^ and 2^nd^ PCs indicated 3-4 hours of heat exposure as a turning point in transcriptional patterns during the prolonged stress response.

**Fig 1.**
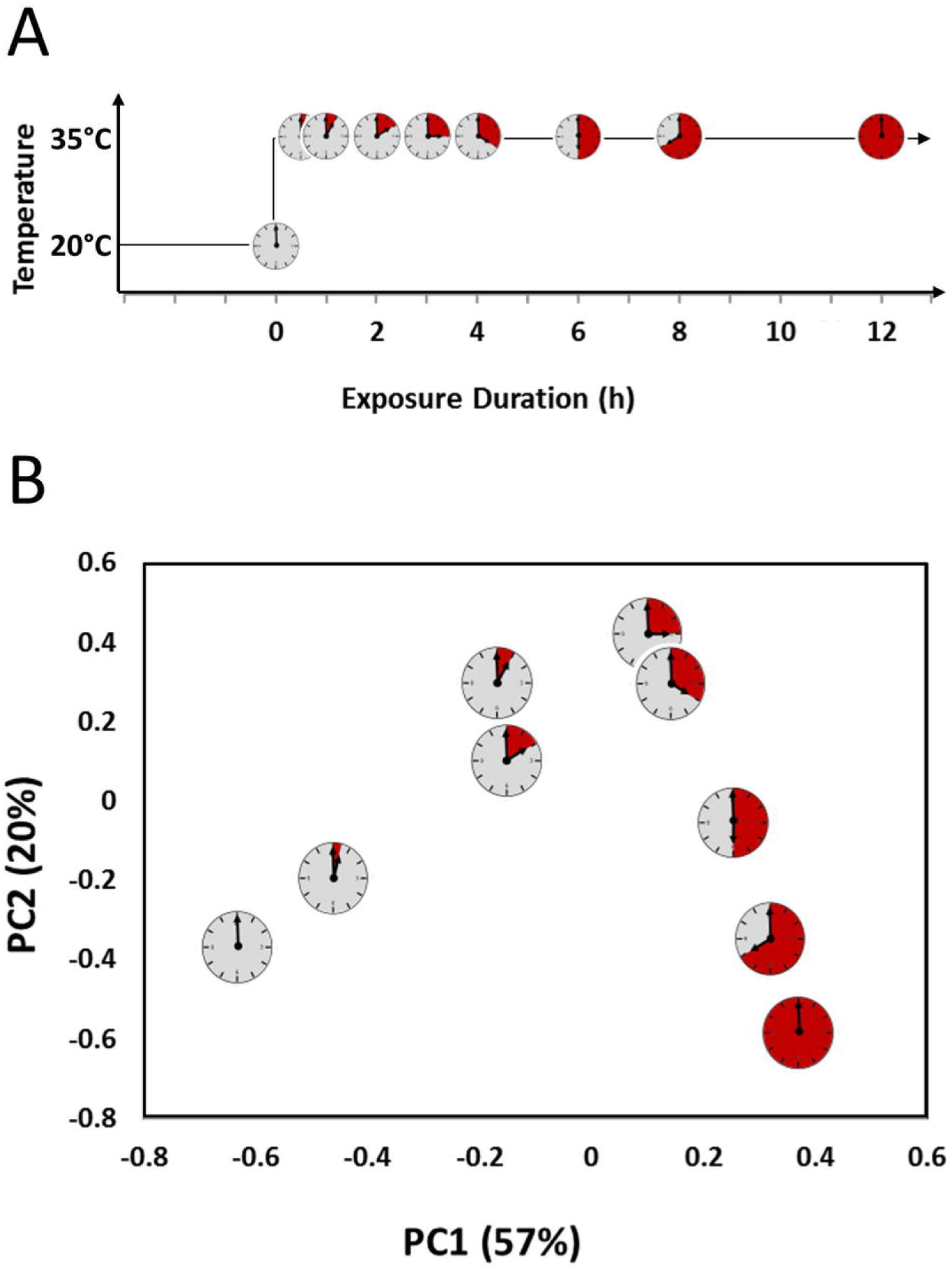
Experimental set-up and principal component analysis. **(A)** mRNA sampling schedule. Bristol N2 populations were grown at 20°C for 46 hours before the start of the heat-shock at 35°C. Clock-symbols indicate the time of sampling for subsequent transcriptome analysis of the dynamic stress response. Each time point (0, 0.5, 1, 2, 3, 4, 6, 8, and 12 hours) was sampled 3-5 times. **(B)** Principal component analysis of gene expression data averaged per time point. The first two components retain 77% of the variation in the data set, and placed the exposure duration (as indicated by the clock symbol) in chronological order.

### Early transcriptional activation of heat shock proteins

Having identified a turning point in transcriptional patterns, we further investigated the temporal dynamics of expression changes for a set of previously associated heat stress response genes. For *C. elegans*, the Gene Ontology database listed 72 genes with a role in the ‘response to heat stress’ (GO:0009408, WormBase version 257). Most of these genes show minor transcriptional changes in the course of the 12-hour heat exposure **(S1 Fig)**. This is not surprising, since many components of the (heat) stress response are constituently expressed [19].

The fastest transcriptional response was found for five heat shock proteins, which are part of the heat shock response pathway activated by HSF-1 upon stress exposure. Within 30 minutes of stress exposure, *hsp-16.1*, *hsp-16.2*, *hsp-16.41*, *hsp-16.48*, and *hsp-70* showed a ∼16-fold increase in expression levels **(Fig S1)**. The expression levels of these genes peak 4 hours into the stress exposure, corresponding with the turning point identified in the PCA. Correlation analysis extracted two more genes from our data set that were not listed in the GO term ‘response to heat stress’, but presented with similar expression patterns to the above heat-shock proteins: F13E9.1 (ortholog of human NISCH) and F44E5.5 (member of the *hsp-70* family). F44E5.5 was also detected in previous studies [18,20]. Together, this small set of seven genes presented the strongest, first and immediate reaction of the transcriptional response to stress.

### Changes in gene expression reach a plateau

We further investigated the temporal dynamics of global transcriptome changes. About ∼6200 (∼30%) genes contributed significantly (q-value < 0.01) to the variation explained by the first two principal components. This sub-set was used as input for k-means clustering to extract common patterns in gene expression changes, identifying six distinct stress-response groups (Fig 2A-B). Cluster 1, 4, and 6 (representing ∼3790 (60%) of the genes) contained genes downregulated during exposure to heat (Fig 2A), and cluster 2, 3, and 5 contained upregulated genes (∼2450 (40%) of genes; Fig 2B). The largest changes were found in cluster 1 and 3 with an average 5-fold down- and 32-fold up-regulation, respectively.

**Fig 2.**
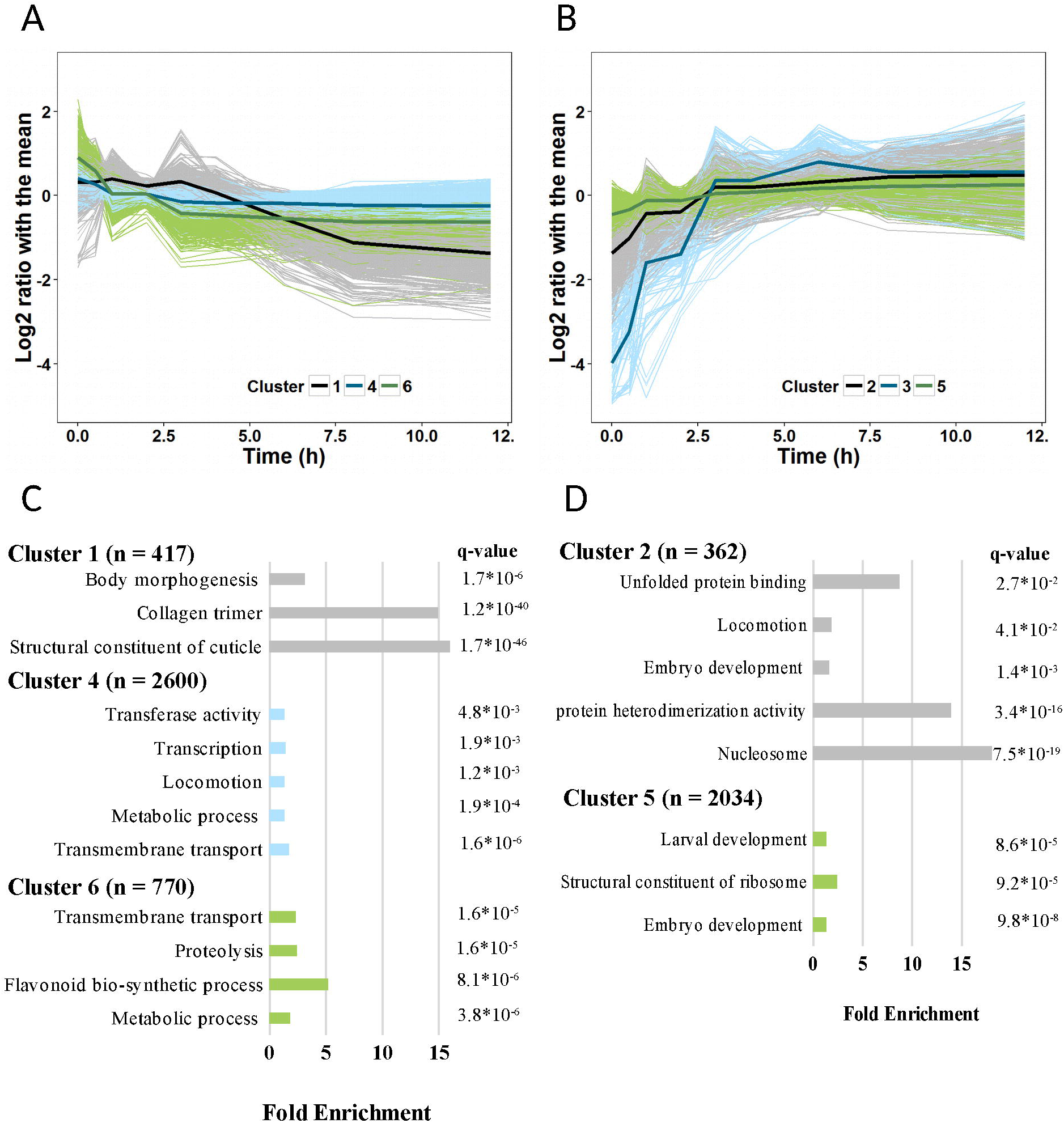
Temporal dynamics and functional enrichments of gene expression in response to continuous heat stress at 35°C. Genes with similar patterns in expression (log2 ratio with the mean) were grouped by k-means clustering. Dark coloured bold lines present the average expression of the individual clusters; lighter corresponding colours present the expression of individual genes. Enrichment analysis of gene clusters was performed with DAVID 6.8. **(A)** Cluster 1, 4, and 6 showed a downward trend in gene expression during heat stress. **(B)** Cluster 2, 3, and 5 were upregulated in response to heat stress. **(C)** Enrichment of downregulated gene clusters. **(D)** Enrichment of upregulated gene clusters 2 and 5. Analysis of cluster 3 did not result in a significant enrichment.

Within these clusters, the initial in- or decrease in transcript levels started rapidly, between 0.5-1 hour after initiation of the heat stress exposure, and reached a plateau before 3 hours into the stress response. One exception is cluster 1, consisting of 420 genes that were down-regulated after 4 hours. Interestingly, this was the only pattern clearly distinguishing the later (>4h) time points. As previously indicated by the PCA, the transcriptional patterns reveal a global change in expression dynamics after approximately 3-4 hours into the stress response, starting with a highly dynamic adaptive phase and ending with a plateau phase of minimal overall changes.

To explore the biological functions associated with the gene sets within the individual response clusters, an GO-enrichment analysis was performed (Fig 2C-D; **S1 Table**). Overall, the down-regulated clusters were enriched with structural constituents of the cuticle, particularly collagens (*col, dpy, rol, sqt*), as well as genes associated with transcription (*nhr*), metabolic processes, and locomotion (Fig 2C). In the upregulated clusters, genes involved in nucleosome assembly (*his*) were found to be overrepresented, as well as those regulating embryo and larval development (Fig 2D). Cluster 3, the smallest group (54 genes), had an immediate and strong reaction to the stress. This cluster could not be associated with an enrichment term. Half of the genes within this cluster have not previously been classified with any GO term yet are very likely involved in the response to heat stress.

### Gene expression dynamics correlated with phenotypic changes

Through transcriptome analysis, we identified a turning point around 3-4 hours into the stress response, separating an initially highly dynamic phase from a later mostly stagnant phase. Next, we tested how these observed transcriptional patterns correlate with the effects of increasing heat stress durations on the phenotypic recovery of the animals. To measure the effects, we observed survival, progeny count, and movement in populations that were allowed to recover at 20°C after different heat stress durations (Fig 3). Since it has previously been shown that it can take three days after the exposure to a transient lethal heat-shock to observe the fatal effects in the survival scores of *C. elegans* [11], we recorded daily phenotypic observations over a four day recovery period following the stress.

**Fig 3.**
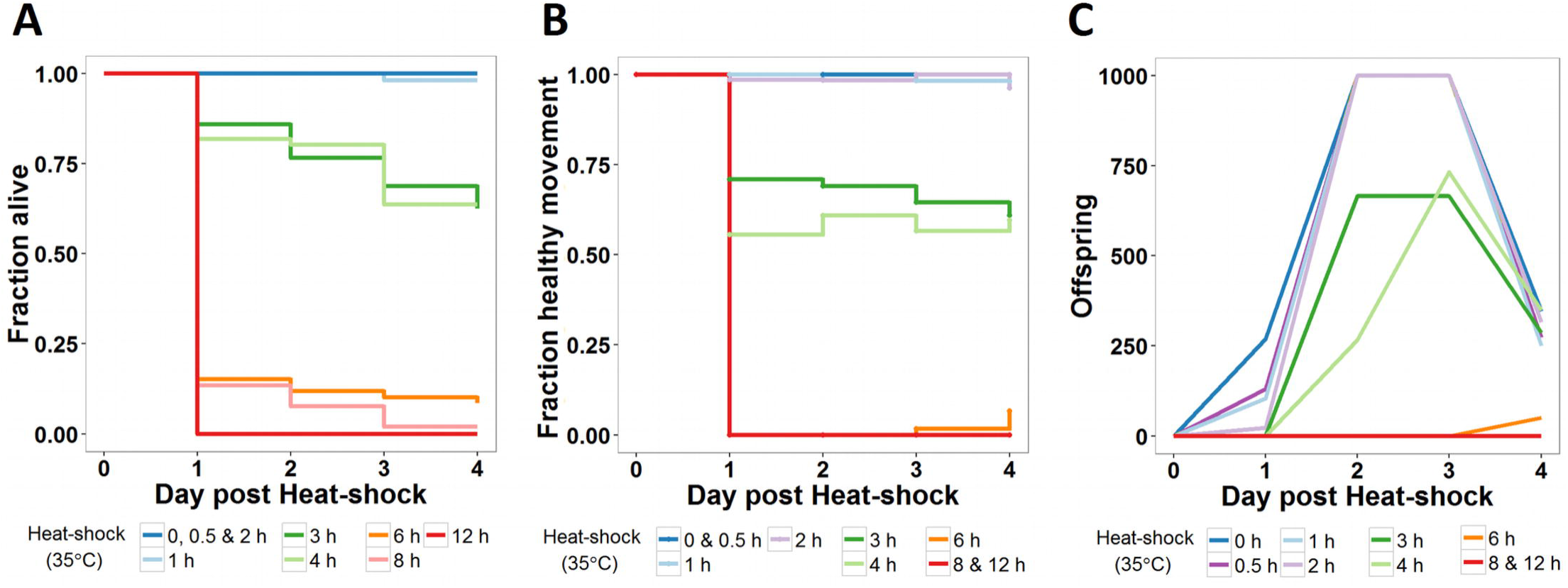
Effect of increasing heat-shock durations on selected phenotypes. After exposing N2 populations in the L4 stage to increasing heat-shock durations at 35°C, populations were maintained at 20°C. A total of ∼65 individuals divided over 3 biological replicas per treatment group were observed. The following phenotypes were scored on the four consecutive days following the heat-shock: **(A)** fraction alive, (**B)** fraction of worms with a healthy movement phenotype (i.e. sinusoidal, constant and unprovoked movement), **(C)** the average number of viable offspring produced per population with a cut-off point set at 1000 offspring.

The heat exposure durations resulted in three phenotypically distinct groups. First, for survival, the animals exposed to heat for up to two hours show high survival chances equal to the control (Fig 3A). An intermediary group was formed by animals exposed to heat for 3-4 hours with about 80% surviving the first day, which steadily declined to ∼60% survival by day 4. It is of note that the exposure duration of this group coincides with the critical time point (3-4h) identified in the transcriptomic data. In the third group, with heat-exposures over 6 hours, survival chances were already drastically reduced after the first day (<20%).

Analogous to the 3 distinct survivorship groups found for short-, intermediate- and long-term stress exposures, this division was also present in the fraction of nematodes regaining a healthy movement during the recovery period, as well as regaining a normal number of progeny (Fig 3B and 3C). The movement in populations exposed to a short heat stress (< 3 hours) did not differ from that of control populations. While the heat stress initially causes slightly lower numbers of progeny, the reproduction peeked 2-3 days after the heat stress together with the control population. For intermediate exposure durations (3-4 hours), 60-70% of animals displayed a normal movement, yet reproduction was further reduced and delayed in these populations. For longer heat exposures (>4 hours), the few surviving individuals commonly presented abnormal, slow, and sporadic movement.

Overall, the transcriptional patterns during heat stress changed dramatically around 3-4 hours which coincided with a phenotypic change, as shown by the drastic decrease in movement, viable offspring, and recovery chances in the days after.

### Heat stress disrupts major developmental processes

To investigate the correlation between gene expression and the different phenotypes, we first looked at how normal developmental processes progressed under heat stress conditions. Snoek *et al.* have dissected the temporal patterns of global transcript levels of *C. elegans* spanning the entire 4^th^ larval stage [21], which corresponds to the time frame used in this study. We analysed the heat stress expression patterns in relation to developmental gene expression. First, we selected gene clusters strongly upregulated during L4 development at 20°C (see Materials and Methods for details; Fig 4A). These genes showed little change in heat stress conditions at 35°C. Likewise, genes with a strong transcriptional response to heat stress (cluster 3) displayed few expression differences during development. Next, we selected gene clusters with a strong decrease in expression levels (Fig 4B). While most of the transcriptional patterns differed between development and heat stress conditions, a relatively small number of genes (*i.e.* 82 genes) were present in both groups. An enrichment analysis of these genes found a strong overrepresentation associated with the cuticle structure and locomotion.

**Fig 4.**
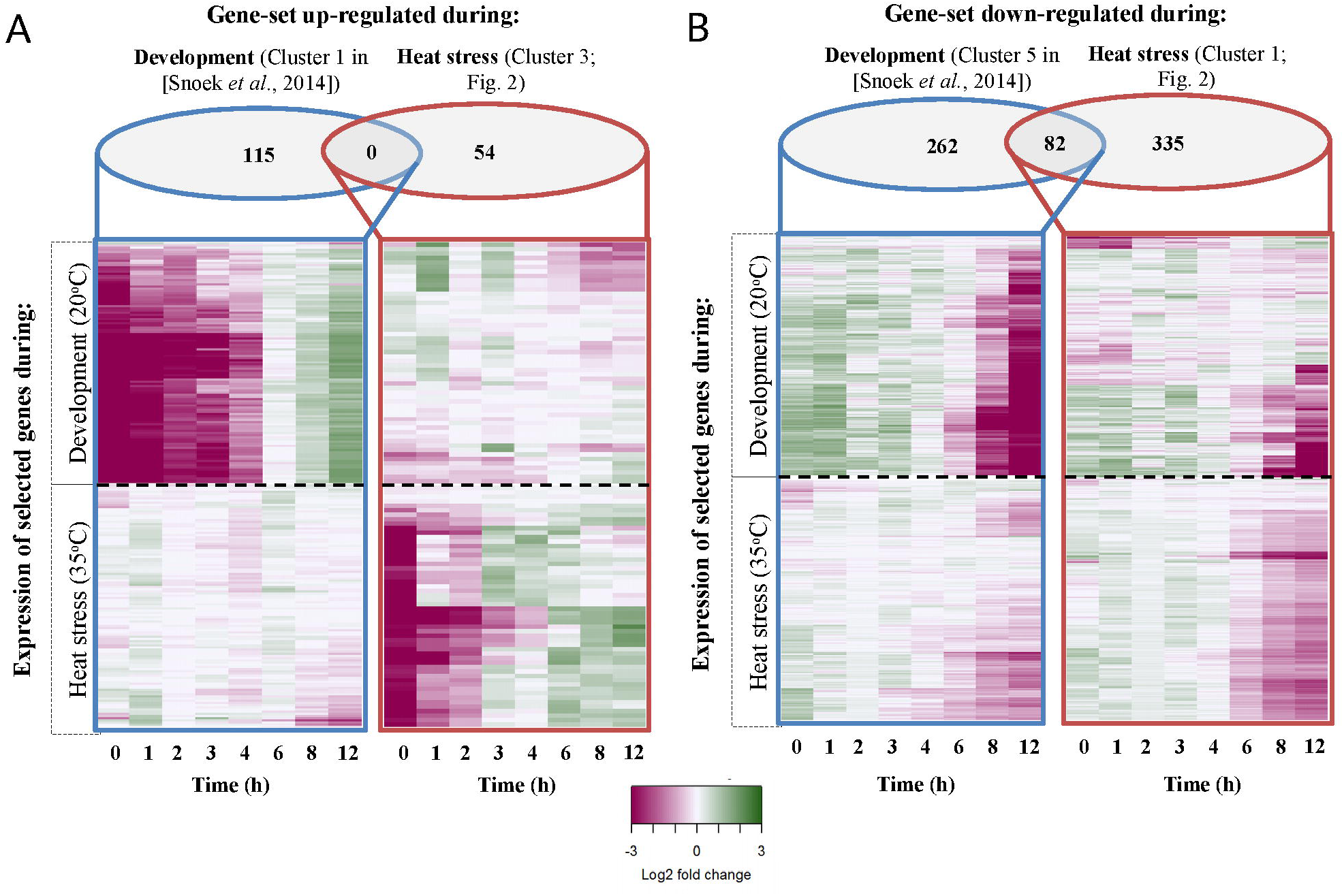
Comparison of expression dynamics during development (20°C; upper panel) and heat stress (35°C; lower panel) based on gene clusters with strong transcriptional patterns. The log2 transformed gene expression is indicated by the colour scale. Developmental gene expression data obtained from [21]. Time was measured beginning 46 hours post age-synchronization. The order of genes within each heat map was retrieved through hierarchical clustering, and is therefore not the same between the upper and lower panels. **(A)** Gene clusters with a strong up-regulation during development (left panel; cluster 1 in [21]) or during heat stress (right panel; cluster 3). **(B)** Gene clusters with a strong down-regulation during development (left panel; cluster 5 in [21]) or during heat stress (right panel; cluster 1). Venn diagrams presents the number of genes within each group.

Together, these results showed that heat stress disrupted the major transcriptional changes normally occurring during L4 development, indicative for the heat stress induced developmental delay. Furthermore, it shows that the animal almost fully shifts its transcriptional program to deal with the acute heat stress conditions.

To identify the genes involved in the sharp decrease in survival after the turning point in expression dynamics, we compared gene expression levels of samples taken between 2-4 hours into the heat-shock with the samples taken at the last three time points (6, 8 and 12 hours). We found 262 upregulated and 667 downregulated genes (q-value < 0.0001; **S1 Table** and **S2 Fig)**. Enrichment analysis revealed an overrepresentation of genes involved with cuticle development and metabolic processes in the late heat stress down-regulated group. Genes involved in reproduction, development, and locomotion were enriched in the late heat stress up-regulated group, possibly showing the continuation of the normal developmental processes after the initial slow down. However, it should be noted that upregulation occurred with very low effect sizes, which could indicate a hampering of the transcriptional processes.

## Discussion

By analysing a series of stress exposure times in *C. elegans* we detected a shift in gene expression patterns between 3-4 hours into the stress response, separating an initially highly dynamic phase from a later mostly stagnant phase. Survival, progeny count, and movement revealed that exposure to a heat stress lasting longer than 4 hours resulted in irreversible damage. Overall, the heat stress response could be divided into three distinct phases: i) an early highly dynamic phase up to 2 hours of stress exposure (including a very early upregulation of heat shock proteins), ii) an intermediate phase in which the transcriptional response attenuates presenting a turning point in dynamics (3-4 hours), iii) a late phase with gradual transcriptomic changes (6-12 hours). Phenotypically, each phase corresponds with distinct trends in the ability to recover from the stress, the ability to recover normal movement, and to produce viable offspring in the four days of recovery following the stress.

To our knowledge, this is the first study that links the dynamics of heat stress response at the transcriptome level to the ability to recover. Gene expression regulation under stress conditions is strictly controlled, its kinetics are rapid and very often it is reversible. This allows for extremely rapid adaptation of cells and tissues in response to general stress, in particular heat stress, and for returning to a baseline level [19]. We analysed the phenotypic recovery from these rapid adaptive changes occurring during stress and found that already a relatively early response to heat stress abruptly changes development.

### Early dynamic response to heat-stress disrupts development

During the early phase of heat stress, the transcriptional response is highly dynamic. About 400 genes (cluster 2 and 5) are highly upregulated. Comparison with transcriptional patterns normally occurring during development has shown that this gene-set specifically reacts in the response to stress. Furthermore, genes highly active during development show low transcriptional changes during stress conditions. These results indicate that the animal almost fully switches its transcriptional focus on counteracting the adverse effects of heat stress. In this context, it is not surprising that heat shock proteins are the first set of genes to show a strong and rapid increase in transcript levels. Shortly after, histones and genes associated with the nucleosome assembly are highly enriched in upregulated gene clusters. Nucleosome remodelling has previously been shown to be an important part of the stress response, *e.g.* by allowing access to transcription sites of stress responsive genes [19,22,23]. In *C. elegans*, depletion of a nucleosome remodelling complex leads to a higher thermal sensitivity [24]. Packaging of DNA into nucleosomes could be an additional protective mechanism during the stress response.

After short exposures to stress, the animals recovered a healthy movement phenotype and started reproducing, indicating that the protective mechanisms put in place by the early transcriptional heat shock response are sufficient in this time frame. However, the disruption of normal transcriptional development could be one of the causes for the observed delay in reproduction. A study in which a two hour 35°C heat-shock was compared to two hour recovery of that heat-shock showed that the transcriptional patterns in the recovery population had still not returned to normal [17]. Also, a delay in reproduction has previously been shown in heat-shocked pre-gravid adult *C. elegans* exposed to temperatures between 30-32°C [15]. Arresting reproduction ensured limited damage to reproductive compartments during stress conditions. We found this delay on a transcriptional level in the early heat stress response, as development and reproduction related genes did not show their normal up regulation.

### Attenuation of dynamic response

At medium-to-long exposures, the transcriptional stress response attenuates corresponding phenotypes, *i.e*. a ∼40% decrease in survival and an increased occurrence of animals with an abnormal movement phenotype. The attenuation of the heat shock response has mostly been studied in several cell lines [24,25]. An integral part is the activation and subsequent suppression of the HSF-1 transcription factor activity through a negative feedback loop, which is partially mediated by those chaperones that are transcriptionally induced by HSF-1, such as HSP-70 [26]. The attenuation of the heat shock response is believed to serve a protective function, as cell lines with defects in the process display lower growth rates and reduced fitness [19,25]. In *C. elegans,* it was shown that a gain-of-function mutation in a negative regulator of the heat-shock response (HSB-1) results in severe effects on survival after heat stress [24]. In our data set, transcripts of chaperones induced by HSF-1 increase immensely within the first 30 minutes of the stress response (**S1 Fig**). The drastic increase slows down until peak levels are reached at 4 hours into the stress response, followed by a small decrease and complete attenuation. It is unclear if the observed global transcriptional slowing down is due to an actively regulated process, such as the HSF-1 feed-back loop, or due to a passive process, such as the accumulation of damage to key cellular processes. Another explanation might be a developmental cue. During normal development without stress, the *C. elegans* transcriptome is highly dynamic, marked by a pronounced shift at 50 hours, which overlaps in time with our point of no return [21]. Passing this point of attenuation might result in the strong decrease in recovery chances. Although the progression of survival rates in the four days following the heat exposure implies that the heat shock does not kill nematodes immediately, the profound damage cannot be repaired.

At long exposures, recovery chances are drastically reduced. While most transcript level have reached a plateau, a distinct exception is the pronounced decrease in expression of a set of genes highly enriched with collagen related genes. Collagens are key components of the nematode cuticle, which is critical for protection and locomotion [27]. During development, the transcription of cuticle collagens is tightly regulated between any of the four molts [27,28]. Comparison with normal development (**S3 Fig**), shows a disruption of these patterns and a general downregulation of all cuticle genes. More recently, gene expression studies in *C. elegans* have shown that collagen genes are highly expressed in short heat stress exposure and during oxidative stress [18,29]. The strongly reduced survival changes after longer exposure might be caused be the later reduction of transcript levels of these cuticle genes in our experiment.

Overall, our study links a strong shift in transcriptional dynamics upon exposure to heat stress with an inability to recover from the stress response. The inability to recover was reflected in a decrease in worm activity, progeny count, and survival in the days after. Therefore, we suggest this critical shift in the dynamics of gene expression marks a point-of-no return ultimately leading to death.

## Materials and Methods

### Nematode Culturing and heat-shock treatment

Hermaphrodites of the *Caenorhabditis elegans* strain Bristol N2 were used for all experiments and kept under standard culturing conditions at 20°C on Nematode Growth Medium (NGM) seeded with *Escherichia coli* strain OP50 as food source [30]. For the experiments, starved populations were placed onto fresh NGM dishes seeded with *E. coli* OP50 by transferring a piece of agar and subsequently grown at 20°C for 3-4 days until sufficient gravid adults had developed. Age-synchronized populations were obtained by bleaching according to standard protocols using a hypochlorite solution [30]. After bleaching, eggs were transferred to fresh 9 cm NGM dishes and maintained at 20°C.

### Heat-shock treatment

The heat shock treatments were performed in an incubator set to 35°C. N2 populations were exposed to the heat stress treatment starting 46 hours after age-synchronization during the L4 stage. We selected the L4 stage because nematodes in this stage exhibit a strong response to heat-stress [11]. The response declines in adult worms [31]. Samples were taken at several time points during the stress period: 0.5, 1, 2, 3, 4, 6, 8, or 12 hours. In total, 3-5 samples were collected for each time point. As preparation for the transcriptome analysis, the populations were washed off the plate with M9 buffer, collected in Eppendorf tubes and flash-frozen in liquid nitrogen and stored at −80°C until further use. For phenotypic observations, the N2 populations were transferred back to pre-heat shock maintenance conditions at 20°C.

## Phenotyping

The selected traits (movement, survival, and progeny count) were observed using a stereomicroscope at approximately 24, 48, 72, and 96 hours post heat-shock. To allow for accurate scoring of all individual animals, the population size per dish was kept at a maximum of 25 animals at the start of the experiment. In total, 3 dishes per heat-shock duration were scored, which amounts to a total of approximately 60 animals per treatment. Animals were transferred to fresh NGM dishes every day during the reproductive phase using a platinum wire. Bagging and suicidal animals were censored.

### Movement and Survival

Movement was scored based on classification systems that have previously been described in association with aging studies, where it acts as a measure of the biological age [32,33]. These systems were combined and adapted to score the impact of the heat-shock. Healthy nematodes are actively moving in a sinusoidal pattern (Hosono: type I; Herndon: Class A). As a result of the heat shock, a proportion of the animals deviated from the healthy phenotype in varying degrees such as visibly lower levels of activity, low responsiveness to touch with a platinum wire and/or an irregular shape of movement (for example due to a partially paralysed tail). This is corresponding to Class B and C of Herndon or Type II and III of Hosono). Worms were scored as dead, when no head movement was observed after 3 touches with a platinum wire.

### Progeny count

It has previously been shown that *C. elegans* can lay non-viable eggs after heat shock [11]. For this reason, the progeny count was measured, defined as the absolute number of living offspring per population. We counted the progeny one day after transferring the adults of the experimental populations to fresh dishes, at which time viable eggs have hatched. For populations with a high level of reproduction, the total number of live offspring was estimated based on the count of a quarter of the dish.

## Transcriptome profile

### RNA isolation

RNA was isolated from the flash frozen samples using the Maxwell® 16 AS2000 instrument with a Maxwell® 16 LEV simplyRNA Tissue Kit (both Promega Corporation, Madison, WI, USA). The mRNA was isolated according to protocol with a modified lysis step. Here, 200 µl homogenization buffer, 200 µl lysis buffer and 10 µl of a 20 mg/ml stock solution of proteinase K were added to each sample. The samples were then incubated for 10 minutes at 65°C and 1000 rpm in a Thermomixer (Eppendorf, Hamburg, Germany). After cooling on ice for 1 minute, the samples were pipetted into the cartridges resuming with the standard protocol.

### Sample preparation and scanning

For cDNA synthesis, labelling and the hybridization reaction, the ‘Two-Color Microarray-Based Gene Expression Analysis; Low Input Quick Amp Labeling’ - protocol, version 6.0 from Agilent (Agilent Technologies, Santa Clara, CA, USA) was followed. The *C. elegans* (V2) Gene Expression Microarray 4X44K slides manufactured by Agilent were used. The microarrays were scanned with an Agilent High Resolution C Scanner using the settings as recommended. Data was extracted with the Agilent Feature Extraction Software (version 10.5) following the manufacturers’ guidelines.

### Data pre-processing

Data was analysed using the ‘R’ statistical programming software (version 3.3.2 64-bit). For normalization, the Limma package was used with the recommended settings for Agilent [34]. Normalization within and between arrays was done with the Loess and Quantile method, respectively [35]. The obtained normalized intensities were log2 transformed and outliers were removed. Batch effects within the data set were calculated with a linear model and removed as previously described [21]. For further analysis, the expression values of biological replicas were averaged. To analyse temporal expression dynamics independent from absolute expression values, the individual intensities measured at each time point for each gene were rescaled to the average expression in time per gene. The obtained values were log2 transformed and are further referred to as the log2 ratio with the mean.

### Data accessibility

The microarray datasets supporting this article have been deposited in the ArrayExpress database at EMBL-EBI (<u>www.ebi.ac.uk/arrayexpress</u>) under accession number E-MTAB-5753.

### Data analysis

Principal component analysis (PCA) was performed on the log2 ratio with the mean to explore the source of underlying variation in gene expression. PCA scores of the first and second component were used to select genes with a significant contribution to the variation in expression dynamics. Selection was based on a significance level p < 0.01 in a linear model relating expression values with PCA scores for the separate components. To explore overall trends in gene expression dynamics, gene clusters were extracted by k-means with 200 iterations on 10 different starting sets. Six clusters were sufficient to visualise distinct patterns in gene expression changes.

Differently expressed genes between time-points were deducted by a linear model using the log2 expression of individual samples (3-5 samples per heat-shock duration). In cases where multiple time points were compared, they were grouped into one factor, e.g. Group 1 (2, 3, 4 hours) vs Group 2 (6, 8, 12 hours). A high significance level of q < 0.0001 was chosen.

To extract genes with similar expression patterns to heat-shock proteins, we used spearman correlation analysis on the log2 ratio with the mean averaged for hsp-16.1, hsp-16.2, and hsp-16.41. Genes were selected with a log2 change >2 within the first 30 minutes of heat exposure.

### Enrichment analysis

To explore the biological functions associated with selected gene sets, we used the functional annotation tool provided by DAVID 6.8 [36,37]. For the enrichment analysis (functional annotation chart), settings were limited to Gene Ontology terms (GOTERM_BP_DIRECT, GOTERM_MF_DIRECT, GOTERM_CC_DIRECT).

### Developmental Data

List of genes within developmental cluster 1 and 5 (strongly up- and downregulated, respectively) were obtained from Snoek *et al.* [21]. The normalized developmental expression data set was retrieved from WormQTL [38,39]. From the developmental time series, a subset of samples were selected that correspond to the heat shock time series (i.e.: of the developmental time series 46h, 47h, 48h, 49h, 50h, 52h, 54h, and 58h corresponding with the time points in the heat shock time series 0h, 1h, 2h, 3h, 4h, 6h, 8h, and 12h, respectively). Expression data of replicates was averaged. To compare expression dynamics between the time series obtained during development and in heat stress conditions, we selected the expression data of subsets of genes with strong expression patterns during development (cluster 1 and 5, Snoek *et al.*, 2014) and heat stress (cluster 1 and 3, Fig 2, **S1 Table**). Heatmaps (R; package: gplots) of the log2 ratio with the mean are used to visualize the comparison of the expression dynamics during development and in heat stress conditions for each subset of genes.

## Supporting information

**S1 Fig.** Temporal expression patterns of heat stress responsive genes during 12 hours of heat stress (35°C). Genes were selected based on the information provided by the Gene Ontology database for the GO term ‘response to heat stress’ (GO:0009408, WormBase version 257). Expression levels of individual genes are presented as the Log2 ratio with the mean.

**S2 Fig.** Volcano plot showing the difference per gene in log2 transformed gene expression levels between medium exposure durations (2, 3, and 4 hours) and long exposure durations (6, 8, and 12 hours). P-values were inferred from a linear model comparing the two groups, and corrected for multiple testing by the Benjamini-Hochberg method. The red line indicates the selected significance level resulting in the selection of 262 upregulated (positive effect size) and 667 downregulated (negative effect size) genes.

**S3 Fig.** Heatmap of temporal expression patterns of structural constituents of the cuticle during development and under heat stress conditions.

**S1 Table.** Gene lists used for GO enrichment analysis, and detailed output of the enrichment analysis performed with the functional annotation tool provided by DAVID 6.8.

